# FIREcaller: Detecting Frequently Interacting Regions from Hi-C Data

**DOI:** 10.1101/619288

**Authors:** Cheynna Crowley, Yuchen Yang, Yunjiang Qiu, Benxia Hu, Armen Abnousi, Jakub Lipiński, Dariusz Plewczyński, Di Wu, Hyejung Won, Bing Ren, Ming Hu, Yun Li

## Abstract

Hi-C experiments have been widely adopted to study chromatin spatial organization, which plays an essential role in genome function. We have recently identified frequently interacting regions (FIREs) and found that they are closely associated with cell-type-specific gene regulation. However, computational tools for detecting FIREs from Hi-C data are still lacking. In this work, we present FIREcaller, a stand-alone, user-friendly R package for detecting FIREs from Hi-C data. FIREcaller takes raw Hi-C contact matrices as input, performs within-sample and cross-sample normalization, and outputs continuous FIRE scores, dichotomous FIREs, and super-FIREs. Applying FIREcaller to Hi-C data from various human tissues, we demonstrate that FIREs and super-FIREs identified, in a tissue-specific manner, are closely related to gene regulation, are enriched for enhancer-promoter (E-P) interactions, tend to overlap with regions exhibiting epigenomic signatures of *cis*-regulatory roles, and aid the interpretation or GWAS variants. The FIREcaller package is implemented in R and freely available at https://yunliweb.its.unc.edu/FIREcaller.

**Highlights:**

– Frequently Interacting Regions (FIREs) can be used to identify tissue and cell-type-specific *cis*-regulatory regions.
– An R software, FIREcaller, has been developed to identify FIREs and clustered FIREs into super-FIREs.

## 1. Introduction

Chromatin folding in the three-dimensional (3D) space is closely related to genome function [1]. In particular, gene regulation is orchestrated by a collection of *cis*-regulatory elements, including promoters, enhancers, insulators, and silencers. Alteration of chromatin spatial organization in the human genome can lead to gene dysregulation and consequently, complex diseases including developmental disorders and cancers [2, 3].

High-throughput chromatin conformation capture (Hi-C) has been widely used to measure genome-wide chromatin spatial organization since first introduced in 2009 [4–6]. Analyzing Hi-C data has led to the discovery of structural readouts at a cascade of resolutions, including A/B compartments [6], topologically associating domains (TADs) [7], chromatin loops [8], and statistically significant long-range chromatin interactions [9–11]. Among these Hi-C readouts identified in mammalian genomes, TADs and chromatin loops are largely conserved across cell types [12, 13], while A/B compartments and long-range chromatin interactions exhibit rather moderate levels of cell-type-specificity [6, 7].

As an attempt to identify Hi-C readouts that are better indicative of cell type or tissue-specific chromatin spatial organizations, we have in our previous work [14], identified thousands of frequently interacting regions (FIREs) by studying a compendium of Hi-C datasets across 14 human primary tissues and 7 cell types. We defined FIREs as genomic regions with significantly higher local chromatin interactions than expected under the null hypothesis of random collisions [14].

FIREs are distinct from previously discovered Hi-C structural readouts such as A/B compartments, TADs, and chromatin loops. In general, FIREs tend to reside at the center of TADs, associate with intra-TAD enhancer-promoter (E-P) interactions, and are contained within broader regions of active chromatin [14]. FIREs are tissue and cell-type-specific, and enriched for tissue-specific enhancers and nearby tissue-specifically expressed genes, suggesting their potential relevance to tissue-specific transcription regulatory programs. FIREs are also conserved between human and mouse. In addition, FIREs have been revealed to occur near cell-identity genes and active enhancers [14]. Thus, FIREs have proven valuable in identifying tissue and cell-type-specific regulatory regions, functionally conserved regions such as enhancers shared by human and mouse, and in interpreting non-coding genetic variants associated with human complex diseases and traits [14–16].

Since the discovery of FIREs, we have collaborated with multiple groups to further demonstrate their value in various applications, resulting in multiple recent preprints and publications [16–19]. For example, in an analysis of adult and fetal cortex Hi-C datasets, FIREs and super-FIREs recapitulated key functions of tissue-specificity, such as neurogenesis in fetal cortex and core neuronal functions in adult cortex [19]. In addition, evolutionary analyses revealed that these brain FIRE regions have stronger evidence for ancient and recent positive selection, less population differentiation, and fewer rare genetic variants [19]. For another example, Gorkin et al. [16] investigated how 3D chromatin conformation in lymphoblastoid cell lines (LCL) varies across 20 individuals. They reported that FIREs are significantly enriched in LCL-specific enhancers, super-enhancers, and immune related biological pathways and disease ontologies, further demonstrating the close relationship between FIREs and *cis-*regulatory elements [16]. In particular, even with the sample size of ≤ 20 individuals, hundreds of FIRE-QTLs (that is, genetic variants associated with the strength of FIRE) have been reported, suggesting that FIREs show strong evidence of genetic regulation.

Despite the importance and utilities of FIREs, only in-house pipelines exist for detecting FIREs, limiting the general application of FIRE analysis and the full exploration of cell-type-specific chromatin spatial organization features from Hi-C data. In this work, we describe FIREcaller, a stand-alone, user-friendly R package for detecting FIREs from Hi-C data, as an implementation of the method described in our previous work [14].

## 2. Materials and Methods

### 2.1 Input matrix

First, FIREcaller takes an *n*×*n* Hi-C contact matrix *M* as the input (Figure 1), which can be from a gzipped text file, or the widely used .hic or .cooler file. The contact matrix *M* is constructed by dividing the genome into consecutive non-overlapping bins of size *b* for each chromosome. In our original work [14], *b* was fixed at 40Kb. In this FIREcaller work, we allow *b* to be 10Kb, 20Kb, or the default 40Kb. Each entry in the contact matrix *M*, *m*_*ij*_, corresponds to the number of reads mapped between bin *i* and bin *j*. The corresponding symmetric *n*×*n* matrix reflects the number of mapped intra-chromosomal reads between each bin pair [6]. We removed all intra-chromosomal contacts within 15Kb to filter out reads due to self-ligation.

**Figure 1.**
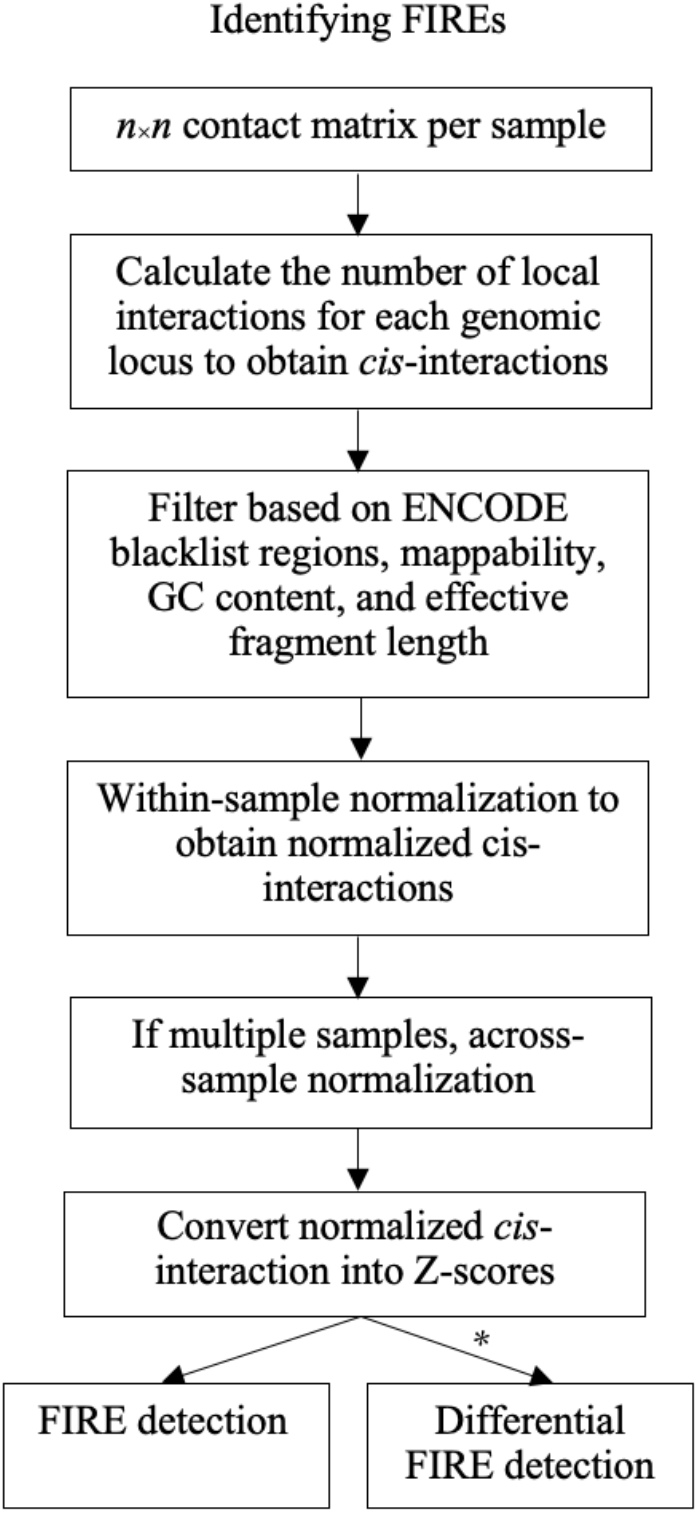
Flow chart of calling FIREs using the FIREcaller software. * indicates when >1 replicate per condition exists. Further detailed in Section 2.8.

Recommendations for the resolution of the input matrix depend on the sequencing depth of the input Hi-C data. Specifically, we recommend using a 10Kb bin resolution for Hi-C data with ~2 billion reads, 20Kb bin resolution for Hi-C data with 0.5-2 billion reads, and a 40Kb bin resolution for Hi-C data with <0.5 billion reads [6, 8, 20–23] (more details can be found in Supplement Information S1).

### 2.2 *Cis*-interaction calculation

Taking the *n*×*n* contact matrix as the input, FIREcaller calculates the total number of local *cis*-interactions for each bin (40Kb default). Following our previous work [14], we define “local” to be within ~200Kb by default. This threshold is largely driven by empirical evidence that contact domains exert influences on transcription regulation within 200Kb. For instance, contact domains reported in human GM12878 cells from *in*-*situ* Hi-C data are at a median size of 185 Kb [8, 20]. In addition, Jin et al. reported a median distance of E-P interactions at 124Kb [21], Song et al. reported ~80% of promoter-interacting regions within 160Kb [24], and Jung et al. found promoter-centered long-range chromatin interactions with a median distance of 158 Kb [25]. Consistently, an analysis of the dorsolateral prefrontal cortex sample [26] showed E-P interactions at a median distance of 157Kb, and our study showed adult cortex E-P interactions at a median distance of 190Kb [19] (Supplement Information S2). On the other hand, multiple *cis*-regulatory regions have been shown to control their target genes from longer genomic distances [3, 19, 20, 27]. To accommodate these longer-range chromatin interactions, our FIREcaller software allows a user-specified upper bound of the *cis*-interacting regions.

### 2.3 Bin level filtering

Bins are then filtered based on multiple criteria that may lead to systematic biases, including effective restriction fragment lengths which measures the density of the restriction enzyme cut sites within each bin, GC content, and sequence uniqueness [28, 29]. FIREcaller removes bins with 0 mappability, 0 GC content or 0 effective fragment length. It also removes bins for which more than 25% of their neighborhood (within 200Kb, by default) bins have 0 mappability, 0 GC content or 0 effective fragment length. In addition, any bins with a mappability less than 90% are removed. Finally, any bins overlapped within the MHC region or the ENCODE blacklist regions [30] are also filtered out (Supplement Information S3).

### 2.4 Within-sample normalization

FIREcaller then uses the HiCNormCis method [14] to conduct within-sample normalization. HiCNormCis adopts a Poisson regression approach, adjusting for the three major sources of systematic biases: effective fragment length determined by restriction enzyme cutting frequency, GC content, and mappability [14].

As a brief summary of the HiCNormCis method, we let *U*_*i*_, *F*_*i*_, *GC*_*i*_ and *M*_*i*_ represent the total *cis*-interactions (15-200Kb, by default), effective fragment length, GC content, and mappability for bin *i*, respectively. We assume that *U*_*i*_ follows a Poisson distribution, with mean *θ*_*i*_, where log(*θ*_*i*_) = *β*_0_ + *β*_*F*_*F*_*i*_ + *β*_*GC*_*GC*_*i*_ + *β*_*M*_*M*_*i*_. After fitting the Poisson regression model, we define the residuals *R*_*i*_ from the Poisson regression as the normalized *cis-*interaction for bin *i* which are approximately normal (Supplement Information S4).

FIREcaller fits a Poisson regression model by default. Users can also fit a negative binomial regression model. In practice, both Poisson regression and negative binomial regression model achieve similar effect of bias removal, while Poisson regression is computationally more efficient (Supplement Information S5).

Our FIREcaller package also allows users to directly input a normalized contact map, for example, data normalized by a different normalization pipeline, via the “normalized” option. By default, normalized=FALSE, if switched to TRUE, FIREcaller will bypass this within-sample normalization step.

### 2.5 Across-sample normalization

If the user provided multiple Hi-C datasets, FIREcaller uses the R function *normalize.quantiles* in the “preprocessCore” package to perform quantile normalization of the normalized *cis*-interactions across samples [31].

### 2.6 Identifying FIREs

FIREcaller then converts the normalized *cis*-interactions into Z-scores, calculates one-sided *p*-values based on the standard normal distribution, and classifies bins with *p*-value < 0.05 as FIREs. The output file contains, for each bin, the normalized *cis*-interactions, the –ln(*p*-value) (i.e., the continuous FIRE score), and the dichotomized FIRE or non-FIRE classification.

### 2.7 Detecting Super-FIREs

FIREcaller also identifies contiguous FIREs, termed as super-FIRE (Figure 2). FIREcaller first concatenates all contiguous FIRE bins by summing their -ln(*p*-value) (i.e., the continuous FIRE score) to quantify the overall or cumulative amount of chromatin interactions. The summed continuous FIRE scores from contiguous FIREs (which we term as super-FIRE score) are then evaluated against their rank from least interactive to most interactive, where FIREcaller determines the inflection point where the slope of the tangent line is one. Super-FIREs are defined as contiguous FIRE regions beyond the inflection point (Figure 2B). This method is adapted from the Ranking of Super-Enhancer (ROSE) algorithm [32], which was originally proposed for the identification of super-enhancers.

**Figure 2.**
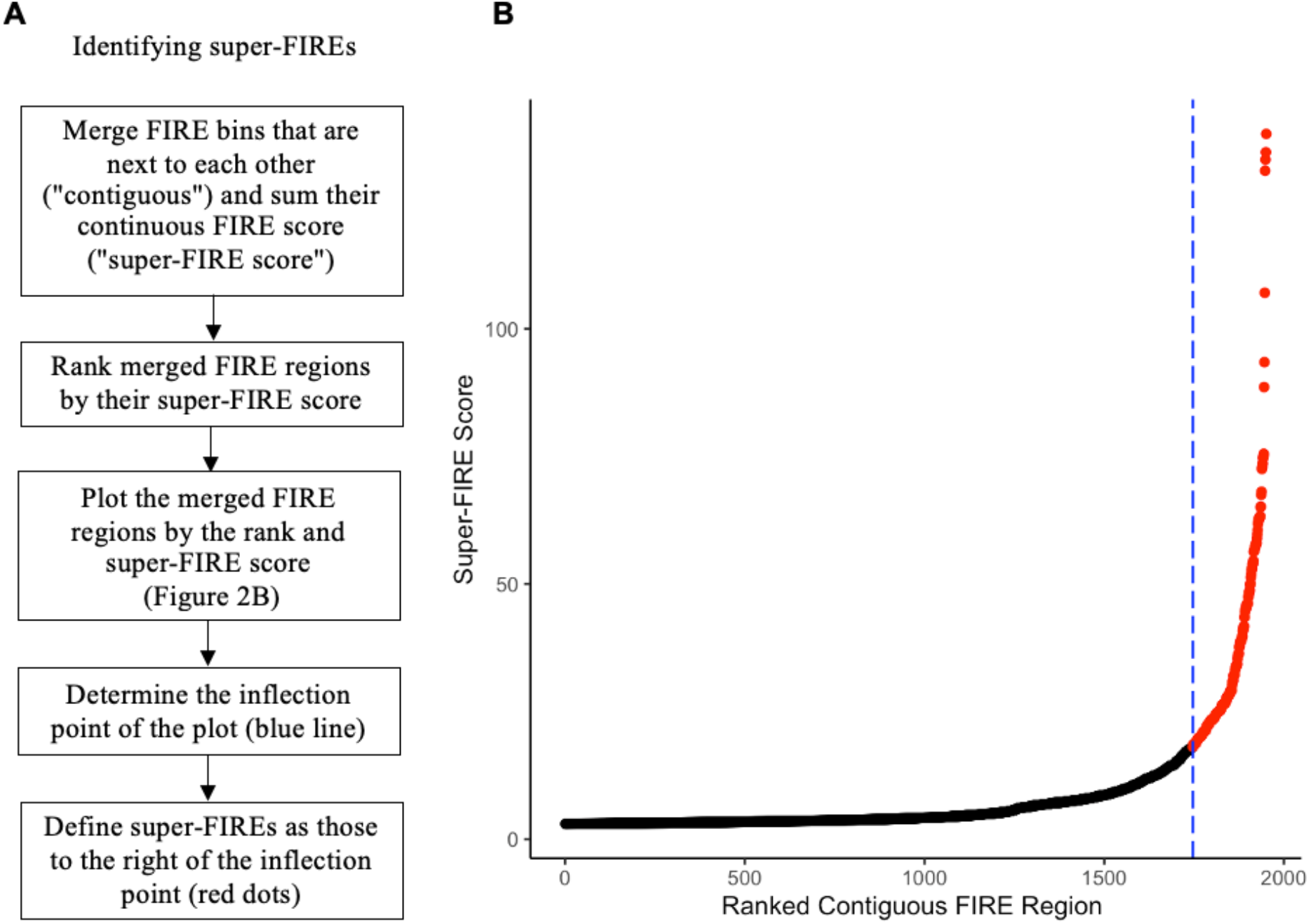
Super-FIRE detection. A) Flow chart for super-FIRE identification. B) Scatterplot of clustered FIREs ranked by their super-FIRE scores for the Hi-C data from hippocampus [14], ordered from the least interactive regions (left) to the most interactive regions (right). Blue dashed line highlights the inflection point of the curve and the red dots highlight super-FIREs, which are clusters of contiguous FIREs to the right of the inflection point.

### 2.8 Identification of differential FIREs

Similar to TADcompare to identify differential TADs [33], FIREcaller allows users to identify differential FIREs between different experimental conditions (e.g., tissues, cell lines, treatments, or developmental stages), when each condition contains at least two replicates. FIREcaller first calculates the normalized *cis*-interactions for each replicate, and then applies the R package “limma” to perform differential FIRE analysis. FIRE bins with fold change > 2 (in terms of the average normalized *cis*-interactions between conditions) and Benjamini-Hochberg adjusted *p*-value < 0.05 are selected as differential FIREs.

### 2.9 Visualizing FIREs and super-FIREs

To visualize FIREs and super-FIREs with other epigenetic data such as TAD boundaries, ChIP-seq peaks and the locations of typical enhancers and super-enhancers, FIREcaller generates a circos plot using the “circlize” package in R [34] (Supplement Information S10).

## 3. Results

To further demonstrate the utility of FIREcaller in terms of connecting the 3D genome structure and function, we first visualized FIREs of Hi-C datasets in Schmitt et al [14] using a virtual 4C plot (Section 3.1) in HUGIn [35], then presented novel FIRE results in fetal [36] and adult brain tissue [37] and integrated with gene expression data (Section 3.2), followed by the joint analysis of E-P interactions, and histone modifications (Section 3.2 – 3.4), as well as differential FIRE analysis (Section 3.5).

### 3.1 An illustrative example

We used the Hi-C data from human hippocampus tissue in our previous study [14] to showcase the utility of FIREcaller. Figure 3 shows an illustrative example of a 400Kb super-FIRE (merged from 10 consecutive bins, and marked by the yellow horizontal bar in the “FIREs” track), which overlaps with two hippocampus super-enhancers (indicated by the two orange horizontal bars in the “Enhancers” track). Notably, this super-FIRE contains a schizophrenia-associated GWAS SNP rs9960767 (black vertical line) [38], and largely overlaps with gene *TCF4* (chr18: 52,889,562-53,332,018; pink horizontal bar depicted at the top with the color of the bar reflecting the log10 expression of the gene), which plays an important role in neurodevelopment [39]. Since rs9960767 resides within a super-FIRE with highly frequent local chromatin interactions, we hypothesize that chromatin spatial organization may play an important role in gene regulation in this region, elucidating potential mechanism by which rs9960767 affects the risk of schizophrenia.

**Figure 3.**
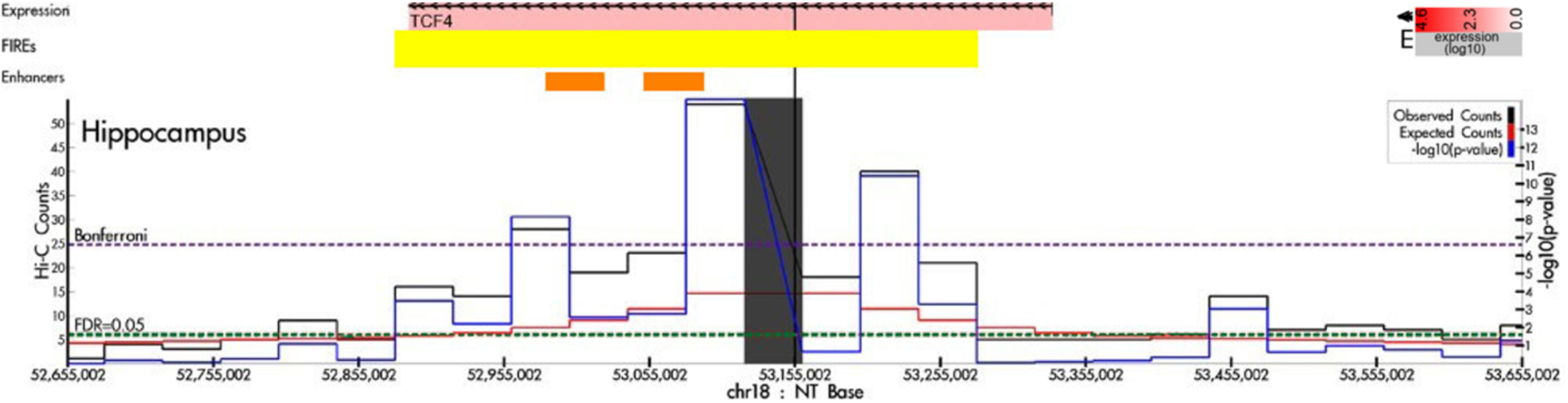
An example of a super-FIRE in human hippocampus tissue. Virtual 4C plot of a 1Mb region (chr18:52,665,002-53,665,002) anchored at the schizophrenia-associated GWAS SNP rs9960767 (black vertical line), visualized by HUGIn [35]. The solid black, red and blue lines represent the observed contact frequency, expected contact frequency, and –log10(*p*-value) from Fit-Hi-C [40], respectively. The dashed purple and green lines represent significant thresholds corresponding to Bonferroni correction and 5% FDR, respectively. The yellow horizontal bar in the “FIREs” track depicts the 400Kb super-FIRE region. The two orange horizontal bars in the “Enhancers” track mark the two hippocampus super-enhancers in the region.

### 3.2 Integrative analysis of FIREs with gene expression in human brain tissues

To study the relationship between FIREs and tissue-specifically expressed genes, we applied FIREcaller to Hi-C data from fetal [36] and adult [37] cortical tissues, and identified 3,925 fetal FIREs and 3,926 adult FIREs. Among them, 2,407 FIREs are fetal-specific and 2,408 FIREs are adult-specific (the remaining 1,518 FIREs are shared).

We then overlapped FIREs with gene promoters and found that the dynamics of FIREs across brain developmental stages are closely associated with gene regulation dynamics during brain development (Figure 4). Specifically, we examined expression levels of genes whose promoters (defined as ± 500 bp of transcription start site [TSS]) overlap with fetal brain-specific FIREs and are expressed in fetal brain, similarly genes whose promoter overlap with adult brain-specific FIREs and are expressed in adult brain. Gene expression data in both fetal and adult brain cortex are from two of our recent studies [36, 37]. These criteria resulted in 707 and 882 genes in fetal and adult brain, respectively. Among them, 412 genes are fetal brain specific, 587 are adult brain specific, and 295 genes are shared (Table 1).

**Table 1.** Tissue-specific FIREs and shared FIREs, and overlapping genes.

|  | # FIREs | # FIREs overlapping with a gene | # of genes overlapping FIREs |
| --- | --- | --- | --- |
| Adult-specific | 2,408 | 488 | 587 |
| Fetal-specific | 2,407 | 338 | 412 |
| Shared | 1,518 | 258 | 295 |

**Figure 4.**
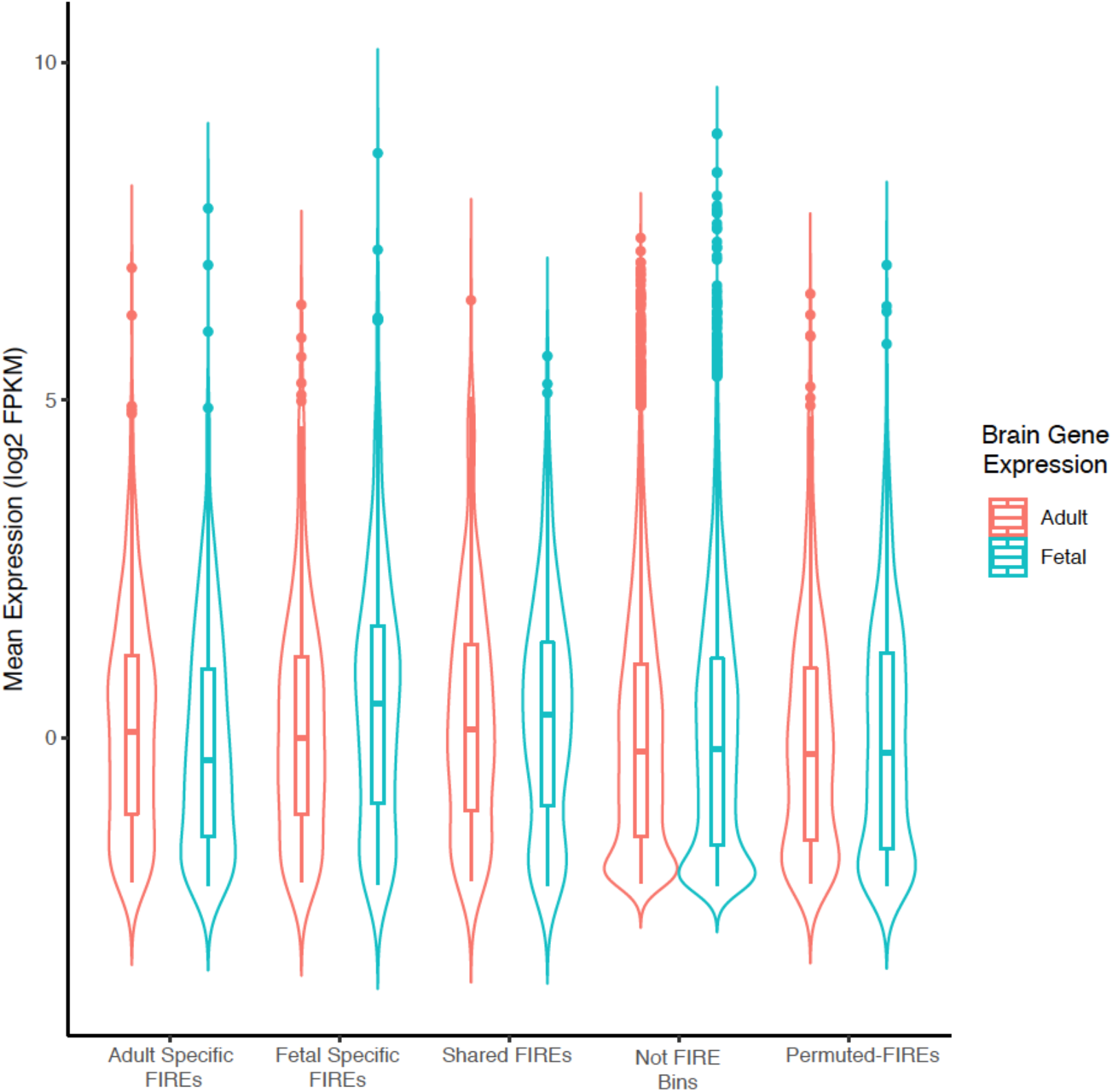
Distribution of expression for genes overlapping fetal or adult brain FIREs. The leftmost pair of violin boxplots shows the expression profile of the 587 genes mapped to adult brain-specific FIREs, with expression measured in fetal brain cortex (blue) and adult brain cortex (red), respectively. The second pair of violin boxplots shows the expression profile of the 412 genes mapped to fetal brain-specific FIREs, again in fetal brain cortex (blue) and adult brain cortex (red), respectively. The third pair shows the expression profile of the 295 genes mapped to FIREs shared between fetal and adult brain, yet again in fetal brain cortex (blue) and in adult brain cortex (red). The fourth pair shows the expression profile of genes not overlapping any FIREs, with a total of 15,640 such genes (labelled “Not FIRE bins”). To the farthest right shows the expression profile of 816 genes overlapping with “permuted-FIREs” with fetal cortex gene expression (blue) and adult brain cortex gene expression (red).

For the 587 genes overlapped adult brain-specific FIREs, the mean gene expression levels, measured by log_2_(FPKM), are −0.052 and 0.190 in fetal and adult brain cortex, respectively. These 587 genes are significantly up-regulated in adult brain (paired *t*-test *p*-value=1.3×10^−10^ Figure 4; Table S5). Meanwhile, for the 412 genes overlapped with fetal brain-specific FIREs, the mean gene expression levels are 0.551 and 0.209 in fetal and adult brain cortex, respectively. These 412 genes are significantly up-regulated in fetal brain (paired *t*-test *p*-value=7.8×10^−13^) (Figure 4; Table S5). By contrast, for the 295 genes overlapped with FIREs shared between fetal and adult cortex, the mean gene expression levels are 0.328 and 0.312 in fetal and adult brain cortex, respectively. These 295 genes show no significant difference in their expression levels between fetal and adult brain (paired *t-*test *p*-value = 0.79). Similarly, genes not overlapping with any FIREs exhibit no significant expression differences in fetal and adult brains either (paired *t*-test *p*-value = 0.96) (Figure 4). For genes overlapped with “permuted-FIREs”, there is no significant difference in expression levels between fetal and adult brain (paired *t*-test *p*-value = 0.84) (Figure 4).

### 3.3 Integrative analysis of FIREs and E-P interactions

We used Hi-C data from left ventricle and liver tissues from Schmitt et al study [14], and applied Fit-Hi-C [40] to call significant chromatin interactions at 40Kb bin resolution. We only considered bin pairs within 2Mb distance. Next, we used H3K27ac ChIP-seq peaks [41] in left ventricle and liver tissues to define active enhancers, and used 500 bp upstream / downstream of TSS to define promoters. A 40Kb bin pair is defined as an E-P interaction if one bin contains a promoter, and the other bin contains an active enhancer. In total, at an FDR<1%, we identified 41,401 and 30,569 E-P interactions in left ventricle and liver, respectively. Among them, 29,096 are left ventricle-specific, and 18,264 liver-specific.

We then applied FIREcaller at 40Kb resolution, and identified 3,643 FIREs in left ventricle and 3,642 FIREs in liver, with 1,186 FIREs shared between these two tissues. We found that FIREs are enriched for E-P interactions compared to non-FIREs for both liver and left ventricle (liver: odds ratio [OR] = 7.2, Fisher’s exact test *p*-value < 2.2×10^−16^; left ventricle: OR = 4.0, *p*-value < 2.2×10^−16^). Comparing between the two tissues, we observed that left ventricle-specific E-P interactions are highly enriched in left ventricle-specific FIREs and liver-specific E-P interactions highly enriched in liver-specific FIREs (OR=3.8, *p*-value <2.2×10^−16^; Table 2). Our results demonstrate that the tissue-specificity of FIREs is closely associated with the tissue-specificity of E-P interactions [14].

**Table 2.**
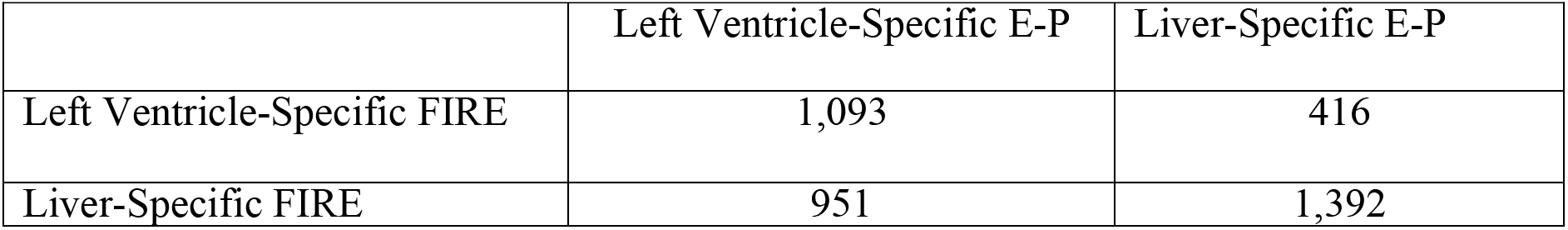
Tissue-Specific FIREs and Tissue-Specific E-P interactions in Liver and Left Ventricle tissues. In the table, we count the numbers of tissue-specific E-P interactions involving tissue-specific FIREs. For example, 1,093 means there are 1,093 left ventricle specific E-P interactions involving left ventricle-specific FIREs. Similarly, for the remaining three counts.

### 3.4 Integrative analysis of FIREs and ChIP-seq peaks

Next, we evaluated the relationship between FIREs and histone modifications in cortex samples [26, 37, 41]. We found that H3K4me3 and H3K27ac ChIP-seq peaks are both enriched at FIRE regions (Figure 5).

**Figure 5.**
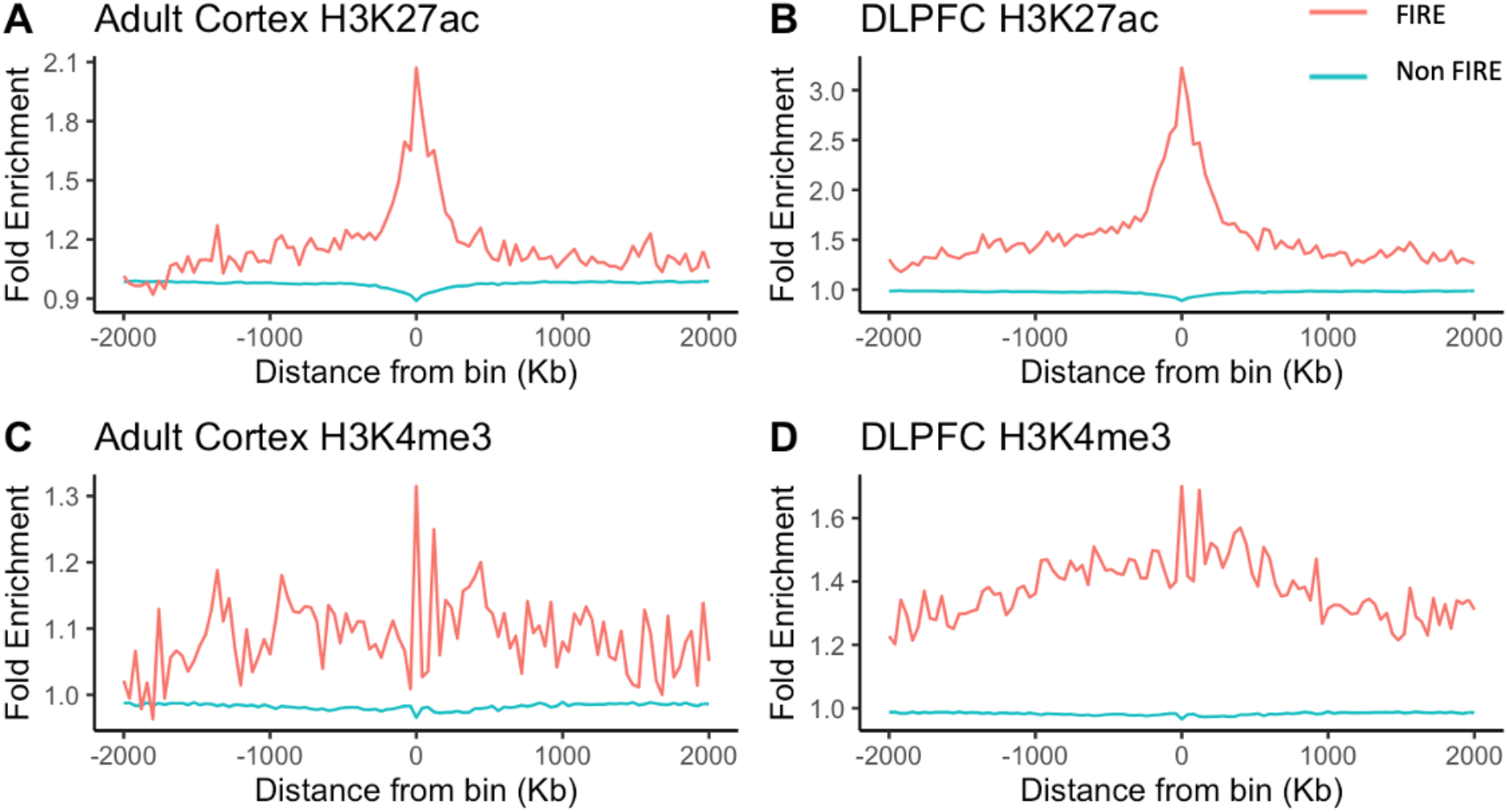
H3K4me3 and H3K27ac ChIP-seq peaks are enriched at FIREs. X axis is the distance from a bin, with the bins grouped into FIRE bins and non-FIRE bins. Y axis is fold enrichment quantified by MACS [42] when applied to the corresponding histone ChIP-seq data.

### 3.5 Differential FIREs between GM12878 and H1 cells

We used FIREcaller to identify differential FIREs between GM12878 cells [8] and H1 embryonic stem cells [14], where Hi-C data for each cell type consists of two biological replicates. We identified 4,140 differential FIREs, where 2,346 FIREs are significantly more interactive in GM12878 and 1,794 more interactive in H1.

Next, we tested whether the differential FIREs are enriched for typical enhancers or super-enhancers [41] in the corresponding cell types. As expected (Figure 6), FIREs more interactive in H1 are significantly more likely to overlap H1 typical enhancers (OR = 1.74; Fisher’s exact test *p*-value = 1.03×10^−4^) and super-enhancers (OR = 1.94; *p*-value = 0.04). Similarly FIREs more interactive in GM12878 are significantly more likely to overlap GM12878 typical enhancers (OR = 78.37; Fisher’s exact test *p*-value < 2.2×10^−16^), and super-enhancers (OR = 78.92; *p*-value < 2.2×10^−16^). We note that the odd ratios for these two cell lines differ rather drastically, which is driven by the fact that H1 FIREs are significantly, but not as strongly enriched in H1 enhancers, compared to GM12878. These results are consistent with those reported in the original Schmitt et al paper [14] where ~35% GM12878 FIREs overlapped with GM12878 typical enhancers, whereas only ~6% H1 FIREs overlapped with H1 typical enhancers (Schmitt et al Figure 4C). Similar patterns were observed for super-enhancers (Schmitt et al Figure 4D).

**Figure 6.**
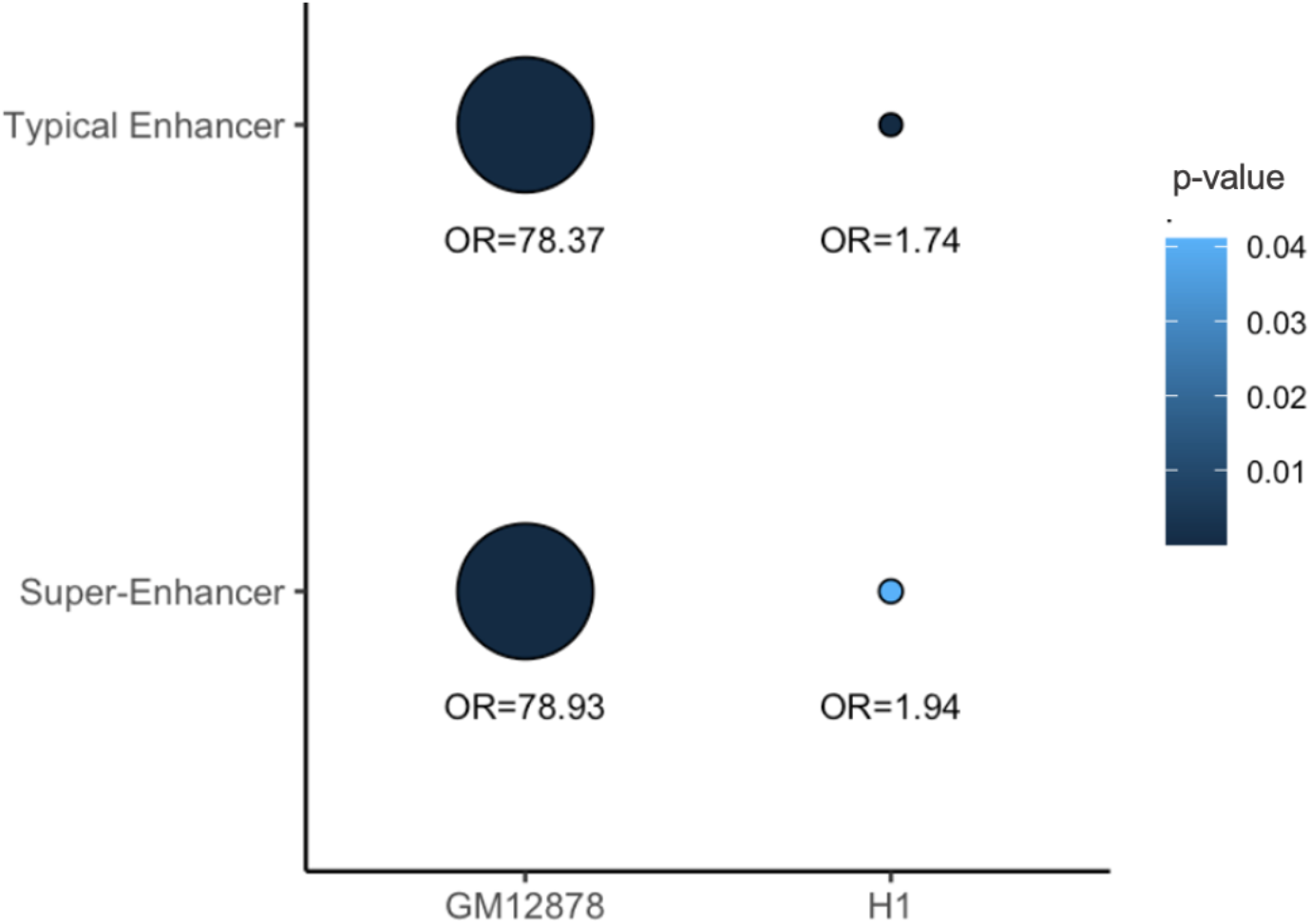
Relationship between differential FIREs and cell-type-specific enhancers in GM12878 and H1 cells. The size of the dots corresponds to the OR and the color of the dots corresponds to the *p*-value.

## 4. Discussion

In this paper, we present FIREcaller, a user-friendly R package to identify FIREs from Hi-C data. We demonstrate its utilities through applications to multiple Hi-C datasets and integrative analyses with E-P interactions, histone modifications and gene expression. We confirmed that FIREs are tissue/cell-type-specific, enriched of tissue/cell-type-specific enhancers, and are near tissue/cell type-specifically expressed genes, informative for prioritizing variants identified from genome-wide association studies (GWAS), consistent with other published works [14, 16–19, 43]. In addition to the identification of FIREs and super-FIREs, our FIREcaller also allows the detection of differential FIREs and visualization of results.

With the development of FIREcaller, FIREs can be easily identified and used in Hi-C data analysis along with TADs, A/B compartments, and chromatin loops. FIREcaller is computationally efficient. Using a single core of a 2.50 GHz Intel processor, the CPU time for running FIREcaller for one Hi-C dataset, all autosomal chromosomes together, at 40Kb resolution with default parameters requires 20.3 seconds with ~113 MB of memory.

Except for the identification of differential FIREs, FIREcaller treats each Hi-C dataset as an independent sample. When multiple replicates are available, user can merge the replicates before calling FIREs and super-FIREs. For differential FIREs, we currently do not allow single replicate as we consider multiple replicates to be necessary for meaningful statistical inference. Future research can explore strategies to accommodate single replicate from each or some conditions. One limitation is that we have not yet applied FIRE calling in many organisms, which warrants future studies. Another limitation of the current FIREcaller is that the resolution might still be too coarse, largely due to the lack of high-depth Hi-C data. With the availability of high-depth Hi-C data based on 4-bp cutters or technologies that allow higher-resolution chromatin architecture mapping in the future, we will explore FIREcaller further at higher resolutions. As a region-based summary of spatial organization information, FIREcaller also lends itself well to sparse data such as those from single cell Hi-C, which warrants further study.

In sum, we developed FIREcaller, a stand-alone, user-friendly R package, to identify FIREs from Hi-C data. We believe FIREcaller is a useful tool in studying tissue/cell-type-specific features of chromatin spatial organization.

## 5. Funding

This research was supported by the National Institute of Health grants R01HL129132 and P50HD103573 (awarded to YL), U54DK107977 and UM1HG011585 (awarded to BR and MH), and R00-MH113823 (awarded to HW). YL is also partially supported by R01GM105785 and U01DA052713. *Conflict of interest*: none declared.

## 6. Authors’ Contributions

**Cheynna Crowley**: Software, Validation, Formal Analysis, Writing – Original Draft. **Yuchen Yang**: Validation, Formal Analysis. **Yunjiang Qiu**: Methodology, Resources. **Benxia Hu**: Formal Analysis, Resources. **Armen Abnousi:** Software. **Jakub Lipinski**: Resources. **Dariusz Plewczynski**: Resources. **Di Wu**: Resources. **Hyejung Won**: Formal Analysis, Resources, Funding Acquisition. **Bing Ren**: Methodology, Funding Acquisition. **Ming Hu**: Supervision, Methodology, Writing- Review & Editing, Funding Acquisition. **Yun Li**: Supervision, Validation, Writing- Review & Editing, Funding Acquisition.

## Notes

https://yunliweb.its.unc.edu/FIREcaller.

### Competing Interest Statement

The authors have declared no competing interest.

## References

1. Dekker, J., M.A. Marti-Renom, and L.A. Mirny, Exploring the three-dimensional organization of genomes: interpreting chromatin interaction data. Nat Rev Genet, 2013. 14(6): p. 390–403.

2. Krijger, P.H. and W. de Laat, Regulation of disease-associated gene expression in the 3D genome. Nat Rev Mol Cell Biol, 2016. 17(12): p. 771–782.

3. Li, Y., M. Hu, and Y. Shen, Gene regulation in the 3D genome. Hum Mol Genet, 2018. 27(R2): p. R228–r233.

4. Tjong, H., et al., Population-based 3D genome structure analysis reveals driving forces in spatial genome organization. Proc Natl Acad Sci U S A, 2016. 113(12): p. E1663–72.

5. Yardimci, G.G., et al., Measuring the reproducibility and quality of Hi-C data. Genome Biol, 2019. 20(1): p. 57.

6. Lieberman-Aiden, E., et al., Comprehensive Mapping of Long-Range Interactions Reveals Folding Principles of the Human Genome. Science, 2009. 326(5950): p. 289.

7. Dixon, J.R., D.U. Gorkin, and B. Ren, Chromatin Domains: the Unit of Chromosome Organization. Molecular cell, 2016. 62(5): p. 668–680.

8. Rao, S.S.P., et al., A three-dimensional map of the human genome at kilobase resolution reveals principles of chromatin looping. Cell, 2014. 159(7): p. 1665–1680.

9. Kaul, A., S. Bhattacharyya, and F. Ay, Identifying statistically significant chromatin contacts from Hi-C data with FitHiC2. Nat Protoc, 2020. 15(3): p. 991–1012.

10. Xu, Z., et al., A hidden Markov random field-based Bayesian method for the detection of long-range chromosomal interactions in Hi-C data. Bioinformatics, 2016. 32(5): p. 650–6.

11. Xu, Z., et al., FastHiC: a fast and accurate algorithm to detect long-range chromosomal interactions from Hi-C data. Bioinformatics, 2016. 32(17): p. 2692–5.

12. Dixon, J.R., et al., Topological Domains in Mammalian Genomes Identified by Analysis of Chromatin Interactions. Nature, 2012. 485(7398): p. 376–380.

13. Rao, Suhas S.P., et al., A 3D Map of the Human Genome at Kilobase Resolution Reveals Principles of Chromatin Looping. Cell, 2014. 159(7): p. 1665–1680.

14. Schmitt, A.D., et al., A Compendium of Chromatin Contact Maps Reveal Spatially Active Regions in the Human Genome. Cell reports, 2016. 17(8): p. 2042–2059.

15. Burgess, D.J., Epigenomics: Deciphering non-coding variation with 3D epigenomics. Nat Rev Genet, 2016. 18(1): p. 4.

16. Gorkin, D.U., et al., Common DNA sequence variation influences 3-dimensional conformation of the human genome. Genome Biol, 2019. 20(1): p. 255.

17. Hu, B., et al., Neuronal and glial 3D chromatin architecture illustrates cellular etiology of brain disorders. bioRxiv, 2020: p. 2020.05.14.096917.

18. Halvorsen, M., et al., Increased burden of ultra-rare structural variants localizing to boundaries of topologically associated domains in schizophrenia. Nat Commun, 2020. 11(1): p. 1842.

19. Giusti-Rodríguez, P., et al., Using three-dimensional regulatory chromatin interactions from adult and fetal cortex to interpret genetic results for psychiatric disorders and cognitive traits. bioRxiv, 2019: p. 406330.

20. Yu, M. and B. Ren, The Three-Dimensional Organization of Mammalian Genomes. Annu Rev Cell Dev Biol, 2017. 33: p. 265–289.

21. Jin, F., et al., A high-resolution map of three-dimensional chromatin interactome in human cells. Nature, 2013. 503(7475): p. 290–294.

22. Schmitt, A.D., M. Hu, and B. Ren, Genome-wide mapping and analysis of chromosome architecture. Nature Reviews Molecular Cell Biology, 2016. 17: p. 743.

23. Lajoie, B.R., J. Dekker, and N. Kaplan, The Hitchhiker’s Guide to Hi-C Analysis: Practical guidelines. Methods (San Diego, Calif.), 2015. 72: p. 65–75.

24. Song, M., et al., Mapping cis-regulatory chromatin contacts in neural cells links neuropsychiatric disorder risk variants to target genes. Nat Genet, 2019. 51(8): p. 1252–1262.

25. Jung, I., et al., A compendium of promoter-centered long-range chromatin interactions in the human genome. Nat Genet, 2019. 51(10): p. 1442–1449.

26. Li, M., et al., Integrative functional genomic analysis of human brain development and neuropsychiatric risks. Science, 2018. 362(6420): p. eaat7615.

27. Fulco, C.P., et al., Systematic mapping of functional enhancer-promoter connections with CRISPR interference. Science, 2016. 354(6313): p. 769–773.

28. Hu, M., et al., HiCNorm: removing biases in Hi-C data via Poisson regression. Bioinformatics, 2012. 28(23): p. 3131–3133.

29. Yaffe, E. and A. Tanay, Probabilistic modeling of Hi-C contact maps eliminates systematic biases to characterize global chromosomal architecture. Nat Genet, 2011. 43(11): p. 1059–65.

30. Amemiya, H.M., A. Kundaje, and A.P. Boyle, The ENCODE Blacklist: Identification of Problematic Regions of the Genome. Scientific Reports, 2019. 9(1): p. 9354.

31. Bolstad, B.M., et al., A comparison of normalization methods for high density oligonucleotide array data based on variance and bias. Bioinformatics, 2003. 19(2): p. 185–93.

32. Whyte, W.A., et al., Master transcription factors and mediator establish super-enhancers at key cell identity genes. Cell, 2013. 153(2): p. 307–319.

33. Cresswell, K.G. and M.G. Dozmorov, TADCompare: An R Package for Differential and Temporal Analysis of Topologically Associated Domains. Front Genet, 2020. 11: p. 158.

34. Gu, Z., et al., circlize Implements and enhances circular visualization in R. Bioinformatics, 2014. 30(19): p. 2811–2.

35. Martin, J.S., et al., HUGIn: Hi-C Unifying Genomic Interrogator. Bioinformatics, 2017.

36. Won, H., et al., Chromosome conformation elucidates regulatory relationships in developing human brain. Nature, 2016. 538(7626): p. 523–527.

37. Wang, D., et al., Comprehensive functional genomic resource and integrative model for the human brain. Science (New York, N.Y.), 2018. 362(6420): p. eaat8464.

38. Stefansson, H., et al., Common variants conferring risk of schizophrenia. Nature, 2009. 460(7256): p. 744–7.

39. Forrest, M.P., et al., The emerging roles of TCF4 in disease and development. Trends Mol Med, 2014. 20(6): p. 322–31.

40. Ay, F., T.L. Bailey, and W.S. Noble, Statistical confidence estimation for Hi-C data reveals regulatory chromatin contacts. Genome research, 2014. 24(6): p. 999–1011.

41. Kundaje, A., et al., Integrative analysis of 111 reference human epigenomes. Nature, 2015. 518(7539): p. 317–330.

42. Zhang, Y., et al., Model-based analysis of ChIP-Seq (MACS). Genome Biol, 2008. 9(9): p. R137.

43. Li, Y., M. Hu, and Y. Shen, Gene regulation in the 3D genome. Human Molecular Genetics, 2018. 27(R2): p. R228–R233.

