## Supplementary file for "FIREcaller: Detecting Frequently Interacting Regions from Hi-C Data"

### Supplementary Information

#### Section S1: Choice of bin resolutions

We tested different bin resolutions using GM12878 Hi-C data [1] with different sequencing depths. We first created 10Kb, 20Kb and 40Kb Hi-C contact matrices using the full data with ~4.4 billion reads. We then down-sampled the full data to 2.2 billion, 1.1 billion and 550 million reads, using a binomial distribution. We repeated this process ten times to generate a total of 20 down-sampled matrices at 2.2 billion reads, 40 down-sampled matrices at 1.1 billion reads, and 80 down-sampled matrices at 550 million reads, at each of the three resolutions (i.e., 10Kb, 20Kb and 40Kb).

Next, we calculated the Jaccard similarity index on the binary FIRE status between all pairs of replicates, at each resolution and sequencing depth. We observed that with ~2.2 billion reads, FIREs identified from each replicate are fairly similar (mean $J$= 0.96 at 40Kb, mean $J$= 0.95 at 20Kb, mean $J$=0.94 at 10Kb resolution; Figure S1 and Table S1). As the resolution increases, the Jaccard similarity index decreases. In general, we recommend using 40Kb resolution for Hi-C data with <1 billion reads, 20Kb for Hi-C data with 1~2 billion reads, and the highest 10Kb resolution only when the Hi-C data has >2 billion reads.

| Resolution | Mean $J$ for 550M | Mean $J$ for 1.1B | Mean $J$ for 2.2B |
|---|---|---|---|
| 10Kb | 0.85 | 0.90 | 0.94 |
| 20Kb | 0.88 | 0.92 | 0.95 |
| 40Kb | 0.90 | 0.94 | 0.96 |

**Table S1. Mean Jaccard similarity index (*J*) for the binary FIRE classifications between replicates for each sequencing depth (~5.5 million reads, ~ 1.1 billion reads, ~2.2 billion reads) at each resolution (10Kb, 20Kb, 40Kb).**

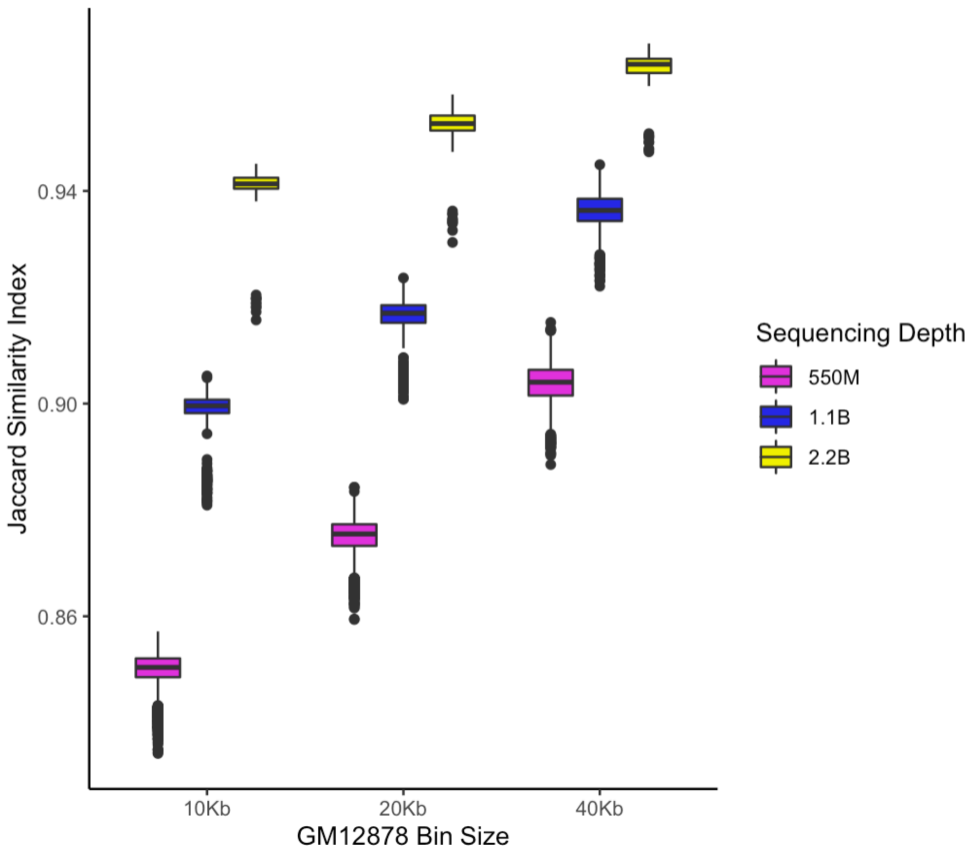


**Figure S1. Boxplots of the Pearson correlations of the dichotomized FIREs for down-sampled GM12878 samples at 10Kb, 20Kb, and 40Kb**, and various sequencing depths of 550M (purple boxplots), 1.1B (blue boxplots), and 2.2B (yellow boxplots) reads.

**Section S2: Distance distribution of enhancer-promoter (E-P) interactions**

Previous studies have shown that the majority of E-P interactions are within 200Kb [1-5]. Here we analyzed high confidence E-P interactions in two brain tissues [6, 7] (Figure S2). We found that high confidence E-P interactions in both adult cortex [6] and in dorsolateral prefrontal cortex (DLPFC) from PsychENCODE [7] have a median distance less than 200Kb (160Kb and 190Kb, respectively; Figure S2).

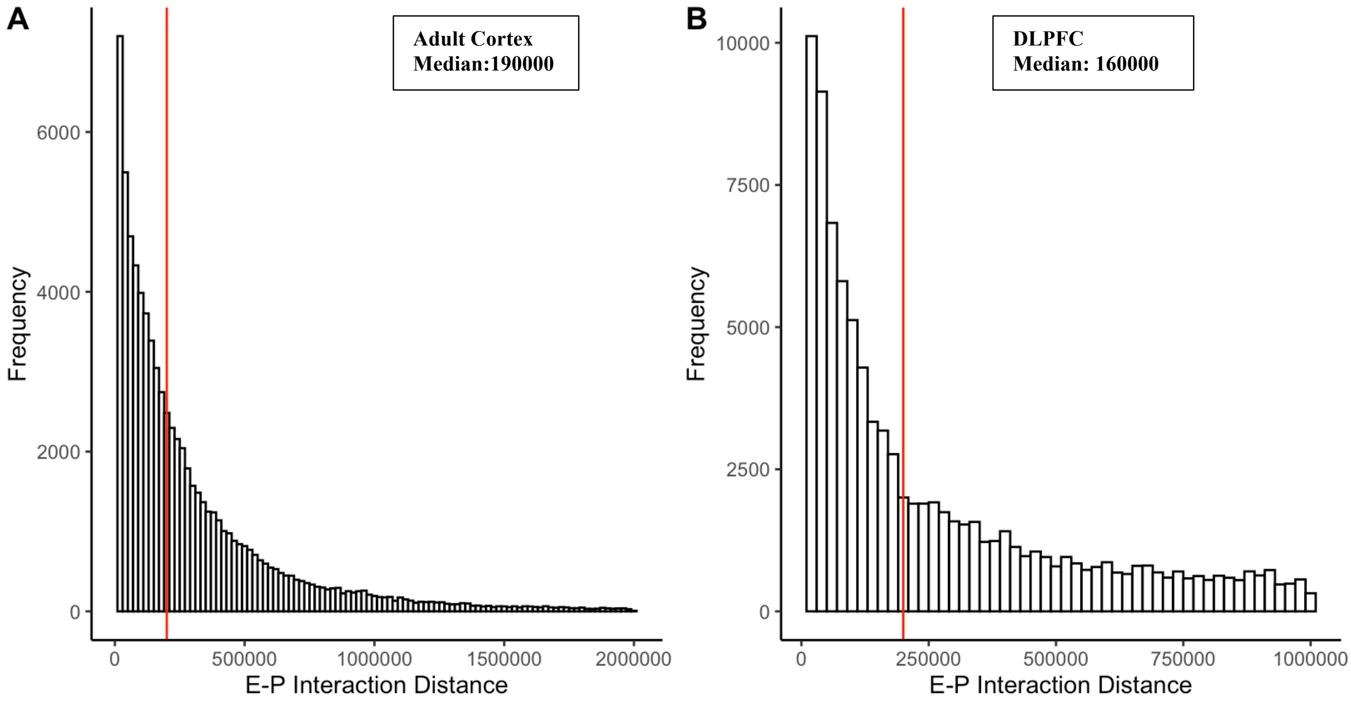


**Figure S2. Histogram of the 1D genomic distance for high confidence E-P interactions. A) Adult Cortex. B) DLPFC**. The vertical red line marks the 200Kb distance.

**Section S3: Bin level filtering**

FIREcaller applies various filtering procedures on the *cis*-interactions to take into account systematic biases and problematic regions. Table S2 lists a breakdown of the number of bins excluded at each filtering step for a Hi-C data with restriction enzyme HindIII with reference genome hg19.

| Bin Size | Total # of bins without filtering | M=0 \| GC=0 \| F=0 Bins | >25% Bad Bins | < 90% M Bins | MHC Region Bins | ENCODE Black List Bins | Total # of bins removed |
|---|---|---|---|---|---|---|---|
| 10Kb | 288,113 | 42,419 (14.7%) | 32,800 (11.4%) | 54,544 (18.9%) | 497 (0.2%) | 1,097 (0.4%) | 61,018 (21.2%) |
| 20Kb | 144,061 | 12,422 (8.6%) | 12,604 (8.7%) | 17,991 (12.5%) | 249 (0.2%) | 602 (0.4%) | 19,149 (13.3%) |
| 40Kb | 72,036 | 5,439 (7.6%) | 6,002 (8.3%) | 7,696 (10.7%) | 125 (0.2%) | 335 (0.5%) | 8,260 (11.5%) |

**Table S2. Number of Bins Removed at each filtering step**. Breakdown of the number of bins at each filtering step by each bin size resolution (10Kb, 20Kb and 40Kb) for an autosomal (chr1-chr22) sample with restriction enzyme Hind3 and genome build hg19. M=Mappability; GC=GC

Content; F=Effective Fragment Length; Bad Bins defined as bins for which more than 25% of their neighborhood bins have 0 mappability, 0 GC content or 0 effective fragment length.

**Section S4: Normality assumption of Poisson regression residuals**

To evaluate the normality assumption of the normalized *cis*-interactions reported from HiCNormCis, we visually inspect a density plot (Figure S3A) and Q-Q plot (Figure 3B). We confirm that the normalized *cis*-interactions are approximately normal, with a slight right skew, justifying the Z-score conversion, and calculating one-sided *p*-values based on the standard normal distribution. These results also justify the normal assumption for the "limma" package [8] used in the differential FIRE analysis.

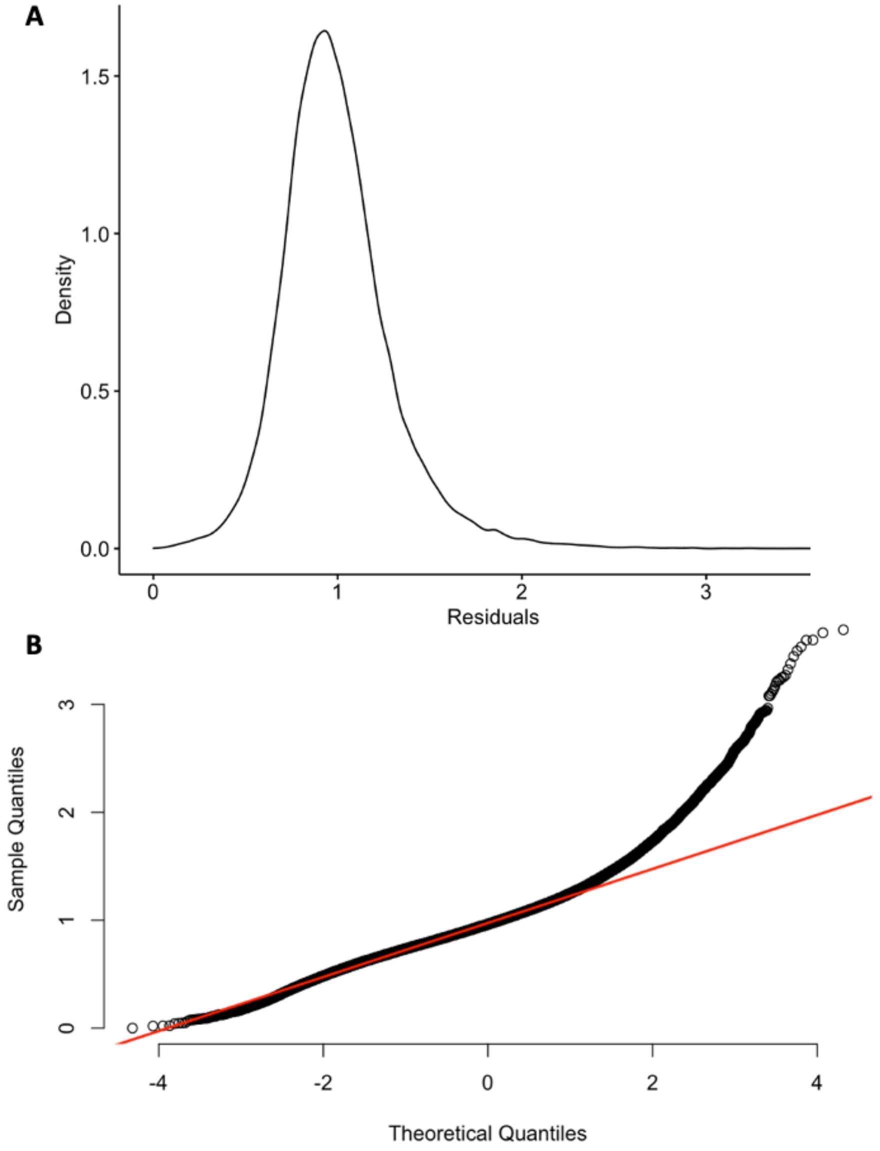


**Figure S3. Normality of Poisson regression residuals (normalized *cis*-interactions) for Hippocampus sample at 40Kb** A) Density Plot B) Q-Q plot

**Section S5: Poisson versus negative binomial models.**

To compare Poisson and negative binomial models, we reported the goodness-of-fit statistics (Table S3), estimation of regression coefficients for systematic biases (Table S4), computational time, and final FIREcaller results (Figure S4). Specifically, we ran FIREcaller using Hi-C data from hippocampus tissue [9], fitting a Poisson regression model and a negative binomial regression model separately.

| | Poisson | Negative Binomial |
|---|---|---|
| Residual Deviance | 536,869 | 65,740 |
| AIC | 944,769 | 608,301 |

**Table S3: Model Goodness-of-Fit Statistics for Poisson Negative Binomial Regression Model.** Smaller residual deviance or AIC indicates a better model fit.

| | Poisson | | Negative Binomial | |
|---|---|---|---|---|
| | Estimate (SE) | *P*-Value | Estimate (SE) | *P*-Value |
| Intercept | -1.19 (0.027) | <0.001 | -1.02 (0.072) | <0.001 |
| Effective Fragment Length | 0.000008 ($1.29\times10^{-7}$) | <0.001 | 0.00009($4.0\times10^{-7}$) | <0.001 |
| GC Content | -0.439(0.0085) | <0.001 | -0.62(0.024) | <0.001 |
| Mappability | 5.12 (0.0267) | <0.001 | 4.93(0.072) | <0.001 |

**Table S4: Regression Estimates for Poisson and Negative Binomial Models**

In terms of model fit, the negative binomial distribution fits the Hi-C count data better (deviance= 65,740) than the Poisson distribution (deviance= 536,869) (Table S3). FIREcaller aims to remove systematic biases and calculate normalized *cis*-contact frequency. We found that these two models resulted in similar estimates of three biases (effective fragment length, GC content and mappability) (Table S4). In addition, these two models result in highly overlapped FIREs. Poisson model and negative binomial model identified 3,642 and 3,760 FIREs, respectively. 3,419 FIREs are shared between two models (odds ratio (OR)= 396.5, Fisher's exact $p$-value< $2.2\times10^{-16}$) (Figure S4).

To compare computational efficiency, we fit the Poisson and the negative binomial regression 100 times. The average running time was 0.44 seconds (0.028 SD) for Poisson regression and 4.28 seconds (0.17 SD) for negative binomial regression.

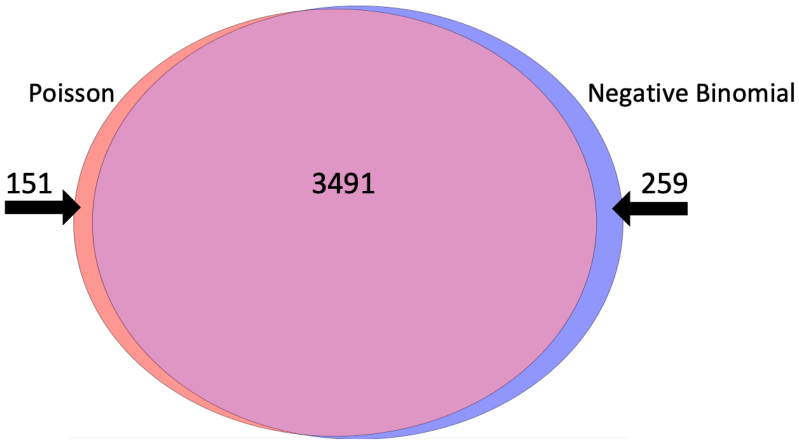


**Figure S4: Venn diagram for FIREs detected from Negative Binomial or Poisson Model**

### Section S6. Evaluation of tissue-specificity.

We applied FIREcaller to Hi-C data from hippocampus, DLPFC, liver, left ventricle, and right ventricle tissues [9], as well as brain tissues from the germinal zone with three biological replicates [10]. We then calculated Pearson correlation ($\rho$) between any pair of samples, based on either the continuous FIRE scores (Figure S5A) or the dichotomized FIRE calls (Figure S5B). As expected, left and right ventricle are highly correlated with each other (continuous FIRE scores: $\rho$=0.89; dichotomized FIREs: $\rho$=0.65), hippocampus and prefrontal cortex are also highly correlated (continuous FIRE scores: $\rho$=0.85; dichotomized FIREs: $\rho$=0.6) and germinal zone brain tissue combined sample (GZ123) is highly correlated with its replicates (continuous FIRE scores:: $\rho$=0.98, 0.99, 0.98; dichotomized FIREs: $\rho$=0.86, 0.91, 0.87 for GZ1,GZ2,and GZ3 respectively).

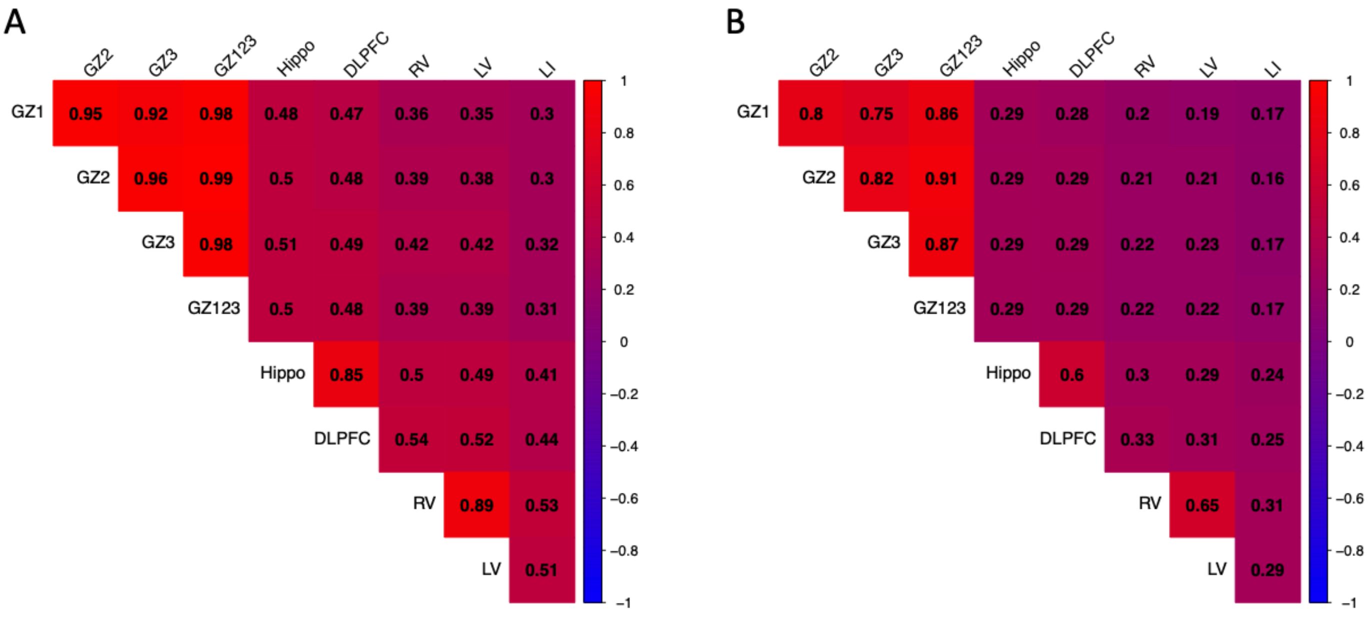

**Figure S5. Pearson correlation of continuous FIRE scores and binary FIREs.** We calculated the Pearson correlations between each pair of samples (one sample each for hippocampus [Hippo], DLPFC, liver [LI], left ventricle [LV], right ventricle[RV], and four samples from germinal zone [GZ] brains: three replicates [GZ1, GZ2, GZ3] and the pooled sample GZ123), for continuous FIRE scores (panel A on the left), or binary FIREs (panel B on the right).

**Section S7: Visualization of Hippocampus tissue raw contact matrix**

To further highlight the region of the 400Kb super-FIRE in human hippocampus tissue, we visually inspected the raw contact matrix (log2(counts)) of both the entire chromosome 18 (Figure S6A) and the ~1Mb region chr18: 52,640,000-53,680,000. Figure S6 shows a higher level of *cis*-interactions (red cluster on heatmap) in the region identified as a super-FIRE (black bar).

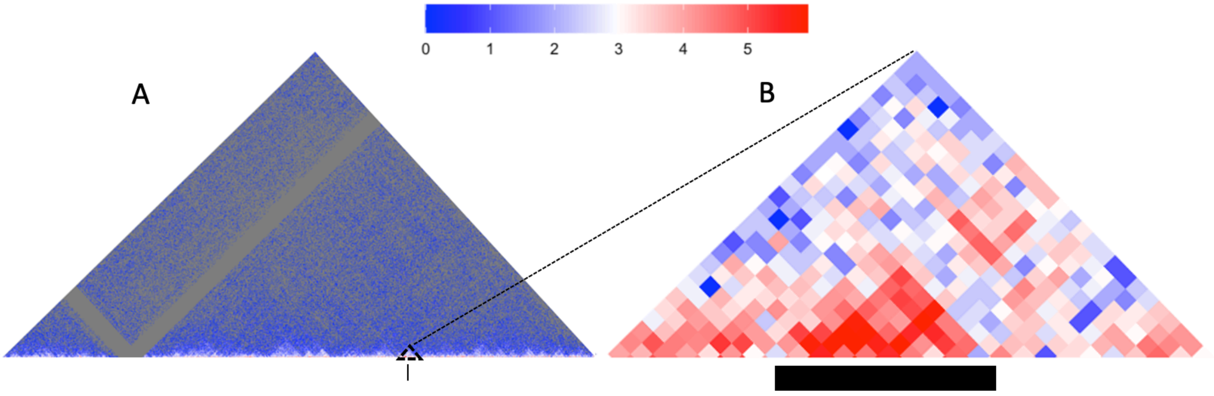


**Figure S6: Heatmap of Hippocampus Tissue logs2 Raw Contact Matrix**. A) A heatmap of the chromosome 18 contact matrix and B) chromosome 18 contact matrix at region 52,640,000, -53680000 with the black bar below showing the 400Kb super-FIRE region.

**Section S8: Comparison of enhancer and promoters in FIREs identified in liver and left ventricle tissue**

To evaluate the enhancer and promoter overlap in FIREs, we first used H3K27ac ChIP-seq peaks [11] in left ventricle and liver tissues to define active enhancers, and used 500 bp upstream / downstream of transcription start site (TSS) to define promoters. We then investigated whether FIREs are enriched for enhancers and promoters. In both tissues, FIREs show significant overlap with enhancers than non-FIREs. Specifically, in liver, OR = 5.22 for enhancers ($p$-value < $2.2\times10^{-16}$), OR = 1.98 for promoters ($p$-value < $2.2\times10^{-16}$); in left ventricle, OR=4.48 for enhancers, ($p$-value < $2.2\times10^{-16}$ ); OR= 1.03 for promoters (with a not significant $p$-value=0.48) (Figure S7).

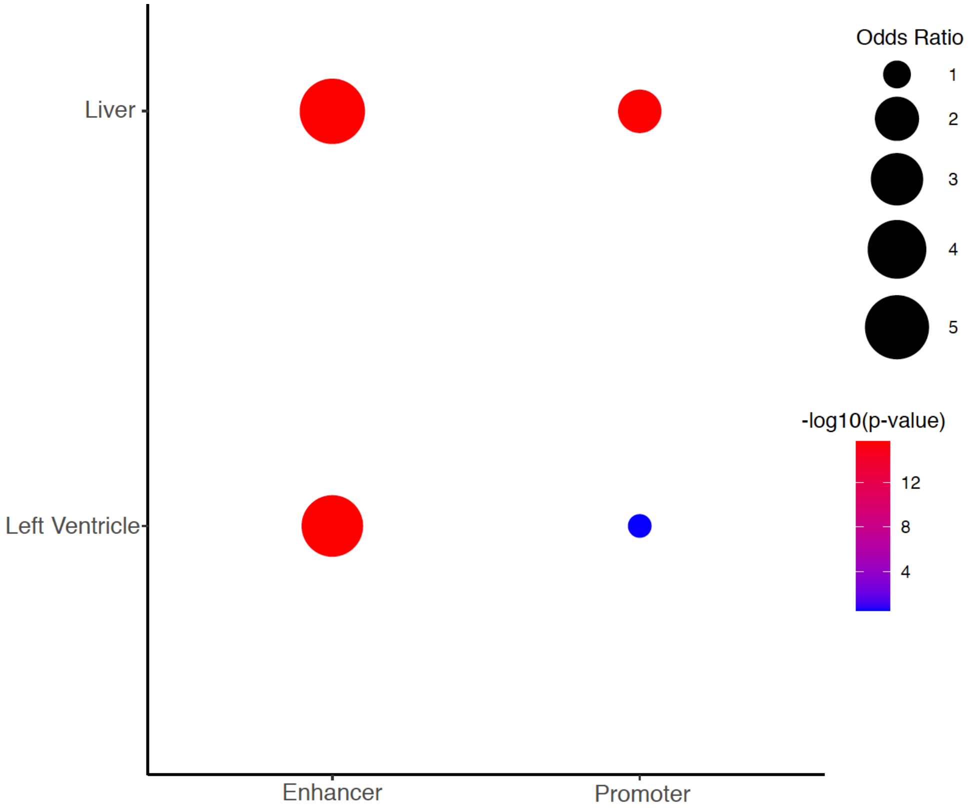


**Figure S7. Dot Plot Showing FIREs Overlapping More with Enhancers than Promoters, in Liver and Left Ventricle tissue.** We use the color scale of blue to red to highlight significance (red being most significant and blue least significant), and the size of the bubble to display the ORs (larger dot indicating larger effect size).

### Section S9: Integrative Analysis of FIREs with Gene Expression in Human Brain Tissue Continued

| | N Genes | Fetal Brain Gene Expression Mean | Adult Brain Gene Expression Mean (SD) | Paired *t*-test |
|---|---|---|---|---|
| Adult-specific FIREs | 587 | -0.052 | 0.190 | *p*-value = $1.256\times10^{-10}$ |
| Fetal-specific FIREs | 412 | 0.551 | 0.209 | *p*-value = $7.79\times10^{-13}$ |
| Shared FIREs | 295 | 0.328 | 0.312 | *p*-value = 0.7854 |

**Table S5. Mean Gene Expression in Adult and Fetal Brain Samples for FIREs and Non-FIREs.**

### Section S10: Visualization Example for Integrating Multi-omic Data

FIREcaller uses the R package "circlize" [12] to visualize FIREs and super-FIREs with other epigenetic data such as TAD boundaries, typical enhancers and super-enhancers. Figure S8 shows FIREs and super-FIREs identified in Hi-C data from hippocampus tissue, chr18, [9] at 40Kb resolution, along with TAD boundaries [9], typical enhancers [11] and super-enhancers [11]. Starting from the outer track and moving inward, first, is an ideogram of chromosome 18; second, the red peaks represent the density of the dichotomized FIREs; third, orange peaks represent the density of the super-FIREs; fourth, the yellow peaks represent the TAD boundaries; fifth, the blue peaks represent typical enhancers; and the most inner lane, purple peaks represent super-enhancers. Note that all underlying features (e.g., FIREs, super-FIREs, TAD boundaries etc) are all binary. The varying height in each circle reflects the density of the binary features.

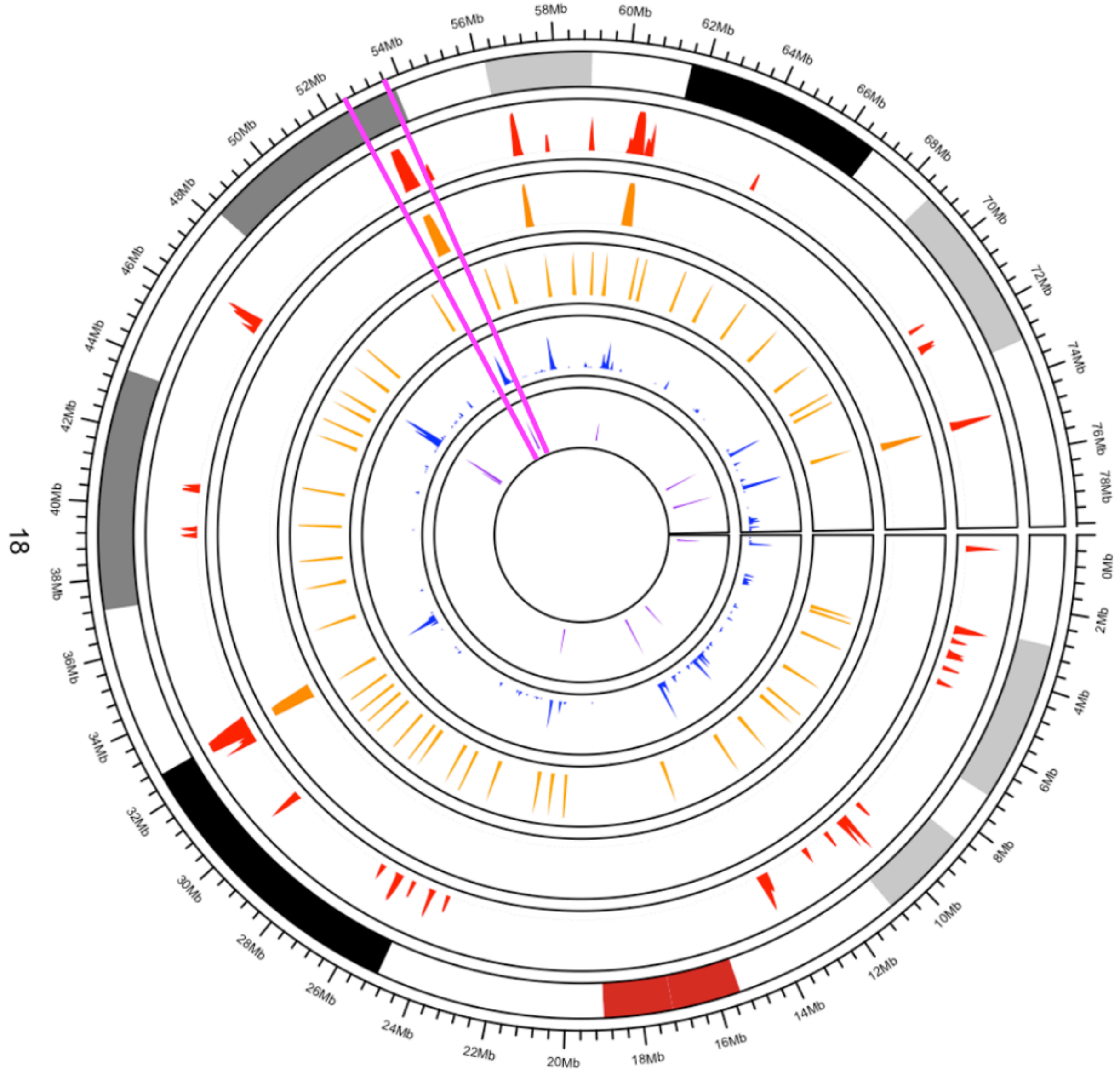


**Figure S8. Circos Plot for multi-omic data for hippocampus Hi-C data highlighting density of the binary FIREs (red), super-FIREs (orange), TAD boundaries (yellow), enhancers**

**(blue), and super-enhancers (purple).** Pink bars highlight the chr18:52,665,002-53,665,002 region visualized in Figure 3 and Figure S6B.

### Section S11: Impact of the Window Size for *Cis*-Interactions

As detailed below, we observed that FIREs detected with the default 200Kb window can be largely captured by larger window sizes. We also noticed that artifacts can occur due to the inclusion of highly interactive regions (e.g., super-FIREs) when increasing the window size, as shown in the example below. Therefore, we urge the user that care needs to be taken when using a larger window size.

First, we compared FIREs called from the hippocampus Hi-C data [9] using the default 200Kb *cis*-interacting window, with FIREs called using a *cis*-interacting window of 300Kb, 400Kb, or 500Kb . We identified 3,642 FIREs using the default 200Kb window size, 3,593 FIREs using a 300Kb window size (80.7 % overlap with FIREs 200Kb window size), 3,563 FIREs using a 400Kb window size (75.3 % overlap with FIREs 200Kb window size), and 3,510 FIREs using a 500Kb window size (70.0 % overlap with FIREs 200Kb window size).

Next, we examined an example where a FIRE was identified at chr1:9,880,000-9,920,000 with a 500Kb window size (FIRE score = 3.07; visualized by the black bar in Figure S9), but was not a FIRE using a 200Kb window size (FIRE score = 0.88), 300Kb window size (FIRE score = 1.82), or 400Kb window size (FIRE score =1.95).

At a 200Kb window size, we identified a super-FIRE spanning chr1:10,080,000-10,480,000, which contributes to the increase of the FIRE scores as the window size increases. More specifically, this region (chr1:9,880,000-9,920,000) identified as a FIRE with a 500Kb window size includes interactions between this region (chr1:9,880,000-9,920,000) and 300Kb of the 400Kb (75%) super-FIRE spanning chr1:10,080,000-10,480,000 (200Kb away from the chr1:9,880,000-9,920,000 region). In contrast, with a 200Kb window, none of the super-FIRE region contributes; with a 300Kb window, 100Kb out of the 400Kb (25%) of the super-FIRE region contributes; and within a 400Kb window, 200Kb out of the 400Kb (50%) of the super-FIRE region contributes.

This example, in our view, represents a FIRE that probably should not be called since the FIRE score with a 500Kb window size does not reflect the property of the region under investigation, but rather a super-FIRE region that happens to be mostly included within the large window size.

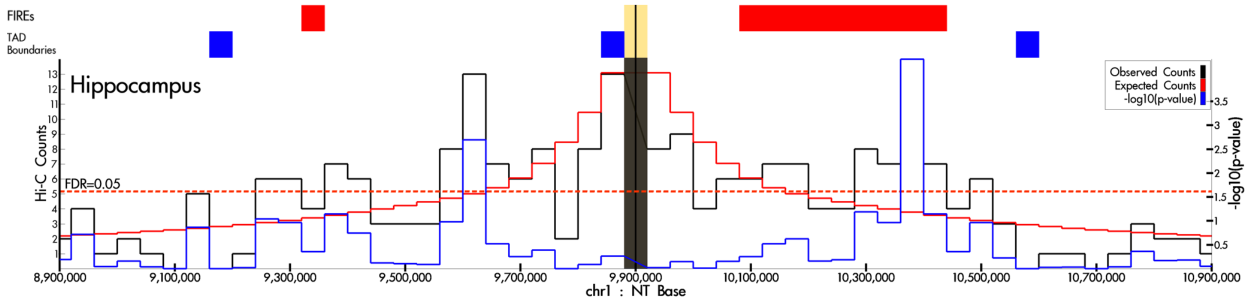

**Figure S9.** Virtual 4C plot of a 2Mb region (chr1:8,900,000-10,900,000) anchored at the 40Kb region identified as a FIRE using a 500Kb *cis*-interacting window (black vertical line), visualized by HUGIn [13]. The solid black, red and blue lines represent the observed contact frequency, expected contact frequency, and –log10(*p*-value) from Fit-Hi-C [14], respectively. The dashed red lines represent significant thresholds corresponding to 5% FDR. The red horizontal bar in the "FIREs" track depicts the FIREs identified using a *cis*-interaction window of 200Kb which includes the 400Kb super-FIRE region. The three blue horizontal bars in the "TAD boundaries" track depicts the TAD boundaries in this 2Mb region.

### Data Availability

All FIREs, super-FIREs and their corresponding input Hi-C contact frequency matrices for human hippocampus tissue have been released at GEO: GSE87112 (PMID: 27851967). In addition, we have made all datasets used in this work, including gene expression data and chromatin marks (H3K4me3 and H3K27ac ChIP-seq peaks), and codes publicly available (website link https://yunliweb.its.unc.edu/FIRE/download.php), to ensure that readers can reproduce our results.

A python pipeline port to the FIREcaller R-package is available at https://github.com/cellular-genomics/python-FIREcaller (contact:).

### Reference


1. Rao, S.S.P., et al., *A three-dimensional map of the human genome at kilobase resolution reveals principles of chromatin looping.* Cell, 2014. **159**(7): p. 1665-1680.
2. Jin, F., et al., *A high-resolution map of three-dimensional chromatin interactome in human cells.* Nature, 2013. **503**(7475): p. 290-294.
3. Yu, M. and B. Ren, *The Three-Dimensional Organization of Mammalian Genomes.* Annu Rev Cell Dev Biol, 2017. **33**: p. 265-289.
4. Song, M., et al., *Mapping cis-regulatory chromatin contacts in neural cells links neuropsychiatric disorder risk variants to target genes.* Nat Genet, 2019. **51**(8): p. 1252-1262.
5. Jung, I., et al., *A compendium of promoter-centered long-range chromatin interactions in the human genome.* Nat Genet, 2019. **51**(10): p. 1442-1449.
6. Giusti-Rodríguez, P., et al., *Using three-dimensional regulatory chromatin interactions from adult and fetal cortex to interpret genetic results for psychiatric disorders and cognitive traits.* bioRxiv, 2019: p. 406330.
7. Li, M., et al., *Integrative functional genomic analysis of human brain development and neuropsychiatric risks.* Science, 2018. **362**(6420): p. eaat7615.
8. Ritchie, M.E., et al., *limma powers differential expression analyses for RNA-sequencing and microarray studies.* Nucleic Acids Res, 2015. **43**(7): p. e47.
9. Schmitt, A.D., et al., *A Compendium of Chromatin Contact Maps Reveal Spatially Active Regions in the Human Genome.* Cell reports, 2016. **17**(8): p. 2042-2059.
10. Wang, D., et al., *Comprehensive functional genomic resource and integrative model for the human brain.* Science (New York, N.Y.), 2018. **362**(6420): p. eaat8464.
11. Kundaje, A., et al., *Integrative analysis of 111 reference human epigenomes.* Nature, 2015. **518**(7539): p. 317-330.

12. Gu, Z., et al., *circlize Implements and enhances circular visualization in R.* Bioinformatics, 2014. **30**(19): p. 2811-2.
13. Martin, J.S., et al., *HUGIn: Hi-C Unifying Genomic Interrogator.* Bioinformatics, 2017.
14. Ay, F., T.L. Bailey, and W.S. Noble, *Statistical confidence estimation for Hi-C data reveals regulatory chromatin contacts.* Genome research, 2014. **24**(6): p. 999-1011.
